# A new pathway for forming acetate and synthesizing ATP during fermentation in bacteria

**DOI:** 10.1101/2020.04.13.039867

**Authors:** Bo Zhang, Courtney Bowman, Timothy J. Hackmann

**Author notes:** Address correspondence to Timothy J. Hackmann.

## Abstract

Many bacteria and other organisms form acetate during fermentation. Forming acetate from high energy-precursors (acetyl-CoA or acetyl phosphate) is one of the few ways that fermentative bacteria generate ATP. Here we found a biochemical pathway for forming acetate and synthesizing ATP that was unknown in bacteria. We found the bacterium *Cutibacterium granulosum* formed acetate during fermentation of glucose. With enzymatic assays, we showed it formed acetate using a pathway involving two enzymes. The first enzyme, succinyl-CoA:acetate CoA-transferase (SCACT), forms acetate from acetyl-CoA. The second enzyme, succinyl-CoA synthetase (SCS), synthesizes ATP. This pathway is common in eukaryotes, but it has not been found in bacteria or other organisms. We found two related bacteria (*C*. *acnes* and *Acidipropionibacterium acidipropionici*) also used this pathway. None used the most common pathway for forming acetate in bacteria (involving acetate kinase and phosphotransacetylase). The SCACT/SCS pathway may be used by many bacteria, not just *C*. *granulosum* and relatives. When we searched genomes for bacteria known to form acetate, we found over 1/6 encoded this pathway. These bacteria belong to 104 different species and subspecies in 12 different phyla. With this discovery, all five pathways known to form acetate and ATP during fermentation can be found in bacteria. This discovery is important to manipulating fermentation and to the evolution of biochemical pathways.

## INTRODUCTION

Acetate is an important product formed during fermentation. Many organisms, from bacteria to multicellular animals, are known to form it (1–4). Its formation is important to energy metabolism of the cell. When the cell forms acetate from high-energy precursors (acetyl-CoA or acetyl phosphate), it can generate ATP by substrate-level phosphorylation (5). This is crucial because fermentation yields few ATP.

The biochemical pathways for forming acetate and ATP during fermentation have been studied for over 80 years (6). Over the course of their study, a total of five pathways have been reported (Fig. 1). For four of the pathways, the high energy precursor is acetyl-CoA. For one pathway, the precursor is either acetyl-CoA or acetyl phosphate.

**Figure 1.**
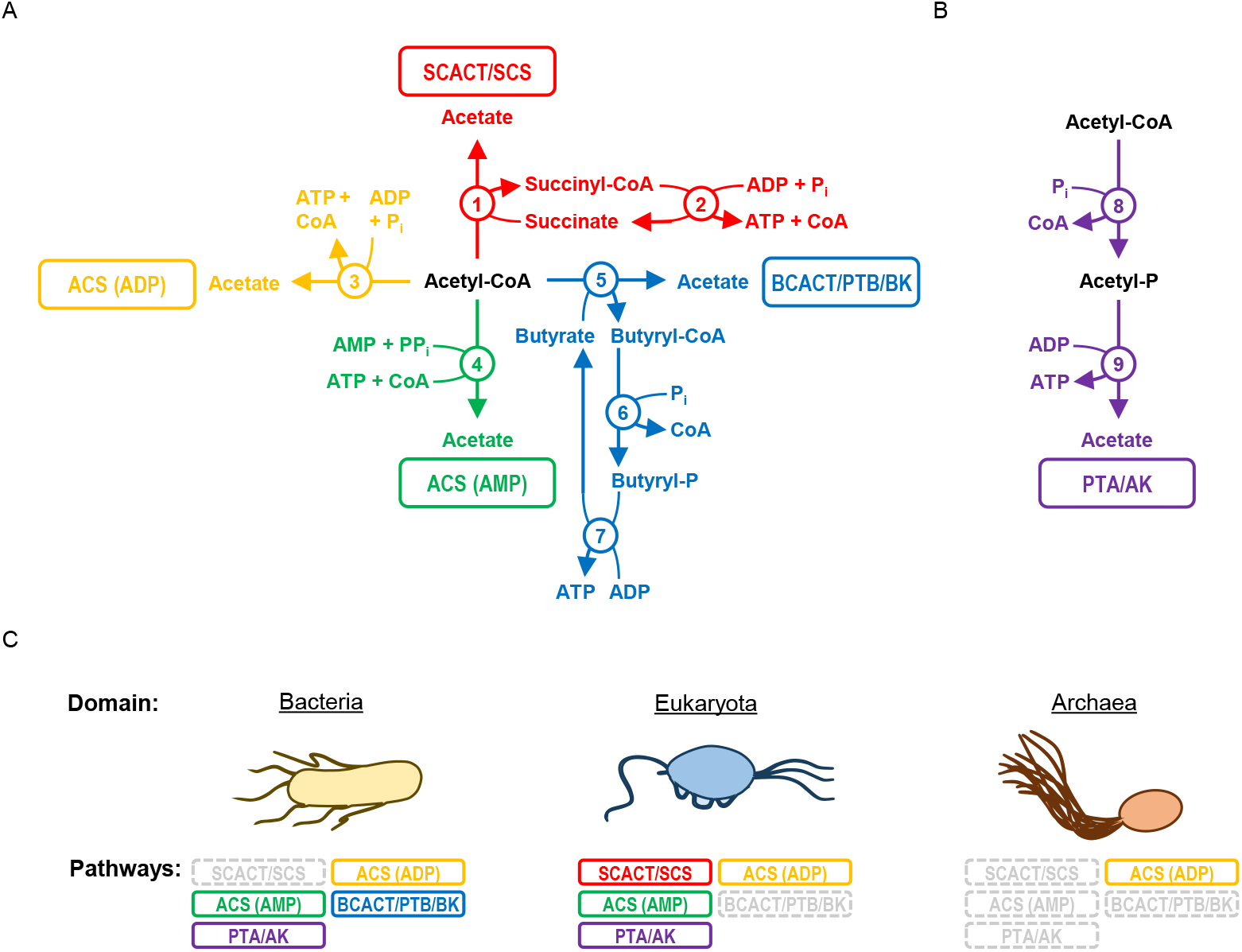
Bacteria use many pathways to form acetate and ATP during fermentation, but none are reported to use the SCACT/SCS pathway. Pathways for forming acetate and ATP from (A) acetyl-CoA and (B) acetyl-CoA or acetyl-P. (C) Reported use of pathways by domain. Pathways that form acetate without ATP (e.g., acetyl-CoA hydrolase; EC 3.1.2.1) are not included. The PTA/AK pathway can use both PTA and AK, or it can use AK alone. Enzymes: 1, succinyl-CoA:acetate CoA-transferase (SCACT; EC 2.8.3.18); 2, succinyl-CoA synthetase (ADP-forming) (SCS; EC 6.2.1.5); 3, acetyl-CoA synthetase (ADP-forming) (ACS [ADP]; EC 6.2.1.13); 4, acetyl-CoA synthetase (ACS [AMP]; EC 6.2.1.1); 5, butyryl-CoA:acetate CoA-transferase (BCACT; EC 2.8.3.8); 6, phosphotransbutyrylase (PTB; EC 2.3.1.19); 7, butyrate kinase (BK; EC 2.7.2.7); 8, phosphotransacetylase (PTA; EC 2.3.1.8); and 9, acetate kinase (AK; EC 2.7.2.1). For bacteria, see ref. (35) for ACS (ADP) pathway, ref. (36) for ACD (AMP) pathway, ref. (37, 38) for BCACT/PTB/BK pathway, and ref. (5) for PTA/AK pathway. For eukaryotes, see ref. (7, 39) for SCACT/SCS pathway, ref. (40, 41) for ACS (ADP) pathway, ref. (42) for ACS (AMP) pathway, and ref. (43, 44) for PTA/AK pathway. For archaea, see ref. (45, 46) for ACD (ADP) pathway. Abbreviations: ACS (ADP), acetyl-CoA synthetase (ADP-forming); ACS (AMP), acetyl-CoA synthetase; AK, acetate kinase; BCACT, butyryl-CoA:acetate CoA-transferase; BK, butyrate kinase; CoA, coenzyme A; P, phosphate; P_i_, inorganic phosphate; PP_i_, pyrophosphate; PTA, phosphotransacetylase; PTB, phosphotransbutyrylase; SCACT, succinyl-CoA:acetate CoA-transferase; and SCS, succinyl-CoA synthetase (ADP-forming).

Curiously, one of the five pathways appears to be missing in bacteria. The pathway missing in bacteria involves the enzymes succinyl-CoA:acetate CoA-transferase (SCACT) and succinyl-CoA synthetase (SCS). It was first reported in a flagellate protozoan (7), and then other eukaryotes (1,8). No report has been made in bacteria (or archaea) (4).

Here we find that the SCACT/SCS pathway is present in bacteria. We find activity of the pathway enzymes in the bacterium *Cutibacterium* (formerly *Propionibacterium*) *granulosum*. We also find activity in certain other propionibacteria. We did not find activity of phosphotransacetylase (PTA) or acetate kinase (AK), which are enzymes of the most common pathway in bacteria (5). From their genome sequences, it appears that over 100 different species and subspecies (type strains) of bacteria encode this pathway to form acetate.

## RESULTS

### C. granulosum forms acetate using the SCACT/SCS pathway

We investigated if *C. granulosum* uses the SCACT/SCS pathway to form acetate because its genome encodes this pathway only. We included other propionibacteria and bacteria for comparison.

To confirm that *C. granulosum* forms acetate during fermentation, we measured its fermentation products when it grew on glucose. We found that *C. granulosum* and other propionibacteria all formed large amounts of acetate (>2.7 mmol/L in culture supernatant) (Fig. 2A).

**Figure 2.**
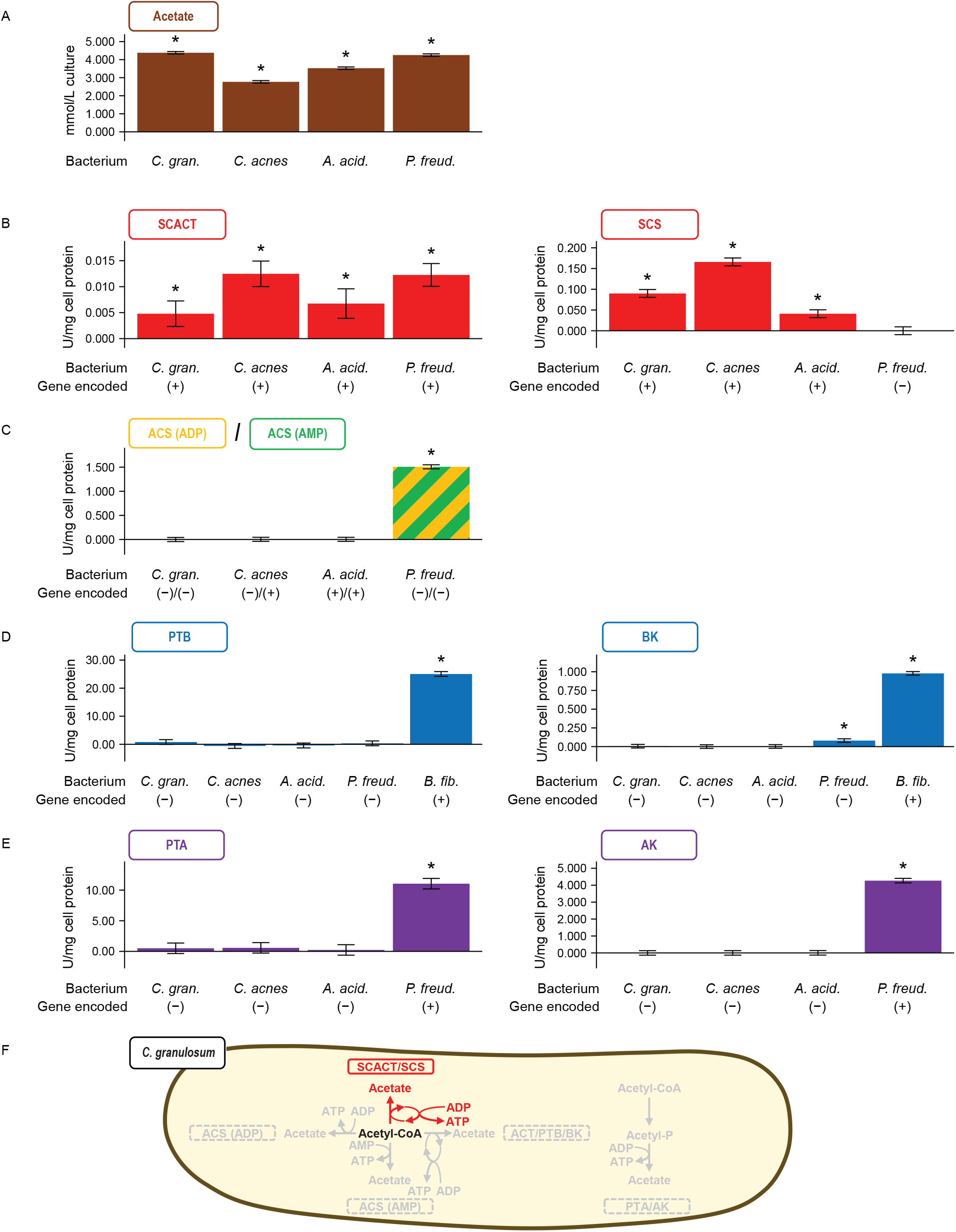
The bacterium *Cutibacterium granulosum* forms acetate using the SCACT/SCS pathway. Other propiniobacteria are included for comparison. (A) Acetate formed during glucose fermentation. (B-E) Activity of enzymes in different pathways for forming acetate. (F) Summary of pathways in *C. granulosum*. In (C), the assay cannot distinguish between the enzymes of ACS (ADP) and ACS (AMP) pathways. In (D), a non-propionibacterium (*Butyrivibrio fibrisolvens*) was included as a control. No attempt was made to measure the activity of BCACT in the BCACT/PTB/BK pathway. Results are means ± standard error of at least 3 biological replicates. Means different from 0 (*P* < 0.05) are marked with an asterisk. Abbreviations: ACS (ADP), acetyl-CoA synthetase (ADP-forming); ACS (AMP), acetyl-CoA synthetase; AK, acetate kinase; BCACT, butyryl-CoA:acetate CoA-transferase; BK, butyrate kinase; PTA, phosphotransacetylase; PTB, phosphotransbutyrylase; SCACT, succinyl-CoA:acetate CoA-transferase; and SCS, succinyl-CoA synthetase (ADP-forming).

To determine if the SCACT/SCS pathway is responsible for forming this acetate, we measured activities of the appropriate enzymes (Fig. 2B–E). We found high activity of both SCACT and SCS in cell extracts of *C. granulosum*. We did not find activity of enzymes in any other pathway. These measurements suggest that *C. granulosum* forms acetate exclusively by the SCACT/SCS pathway. Similar measurements suggest two other propionibacteria (*C*. *acnes*, *Acidipropionibacterium acidipropionici*) use this same pathway, whereas a third (*Propionibacterium freudenreichii*) does not.

To support the accuracy of these measurements, we included multiple controls. When measuring activity of each enzyme, we included one or more bacteria that displayed activity. This supports that our experimental conditions were appropriate to detect activity—i.e., we would have detected activity were any present. Additionally, we spiked cell extracts with purified enzyme at the end of experiments (see Methods). We observed activity after spiking (data not shown). This again supports that our conditions were appropriate to detect activity.

We further supported our enzymatic measurements by analyzing which genes each bacterium encoded (Fig. 2B–E, Table S1). In general, bacteria displayed high activity only when they encoded the appropriate genes. One exception was for acetyl-CoA synthetase (ACS). Two bacteria encoded ACS (ADP-forming) or ACS (AMP-forming), and yet did not display activity. We spiked cell extracts with purified ACS (AMP-forming) and observed activity, confirming that assay conditions were appropriate. Additionally, one bacterium displayed activity even though it did not encode either ACS (ADP-forming) or ACS (AMP-forming). This is due its high activity of AK and PTA; our assay cannot distinguish between ACS and these two enzymes. Together, these results support that *C. granulosum* and certain other propionibacteria form acetate by the SCACT/SCS pathway.

We performed one last experiment to determine if *C*. *granulosum* forms acetate through the SCACT/SCS pathway. If *C*. *granulosum* uses this pathway, it should form acetyl-CoA from acetate, succinate, ATP, and CoA. Indeed, we found cell extracts formed acetyl-CoA from these four substrates (Fig. 3). No acetyl-CoA was formed if any of the four substrates was missing, as expected. In sum, enzymatic and genomic evidence supports that *C*. *granulosum* forms acetate through the SCACT/SCS pathway.

**Figure 3.**
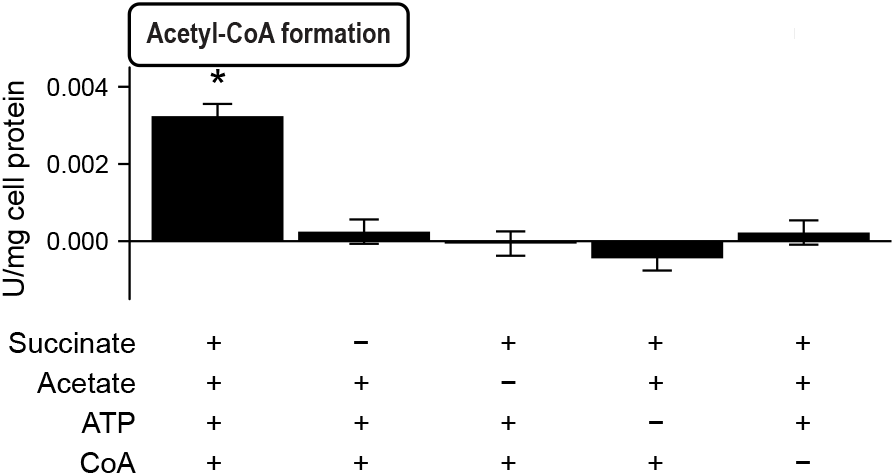
The bacterium *Cutibacterium granulosum* forms acetyl-CoA when provided all four substrates of the SCACT/SCS pathway. In different treatments, substrates were provided (+) or withheld (−). Results are means ± standard error of 4 biological replicates. Means different from 0 (*P* < 0.05) are marked with an asterisk. Abbreviations: CoA, coenzyme A; SCACT, succinyl-CoA:acetate CoA-transferase; and SCS, succinyl-CoA synthetase (ADP-forming).

### Several bacteria encode the SCACT/SCS pathway in their genome

We found that *C. granulosum* encodes the SCACT/SCS pathway by performing a preliminary search of bacterial genomes. To determine how many other bacteria encode this pathway, we performed a systematic search of 2,925 genomes. These genomes represent all species in Bergey’s Manual (9) that are available for analysis on IMG/M (2).

With this search, we found 104 species that 1) encode genes of the SCACT/SCS pathway and 2) are reported to form acetate during fermentation (Fig. 4, Table S1). These species were found across 12 different phyla. This suggests that many different bacteria could use the SCACT/SCS pathway to form acetate.

**Figure 4.**
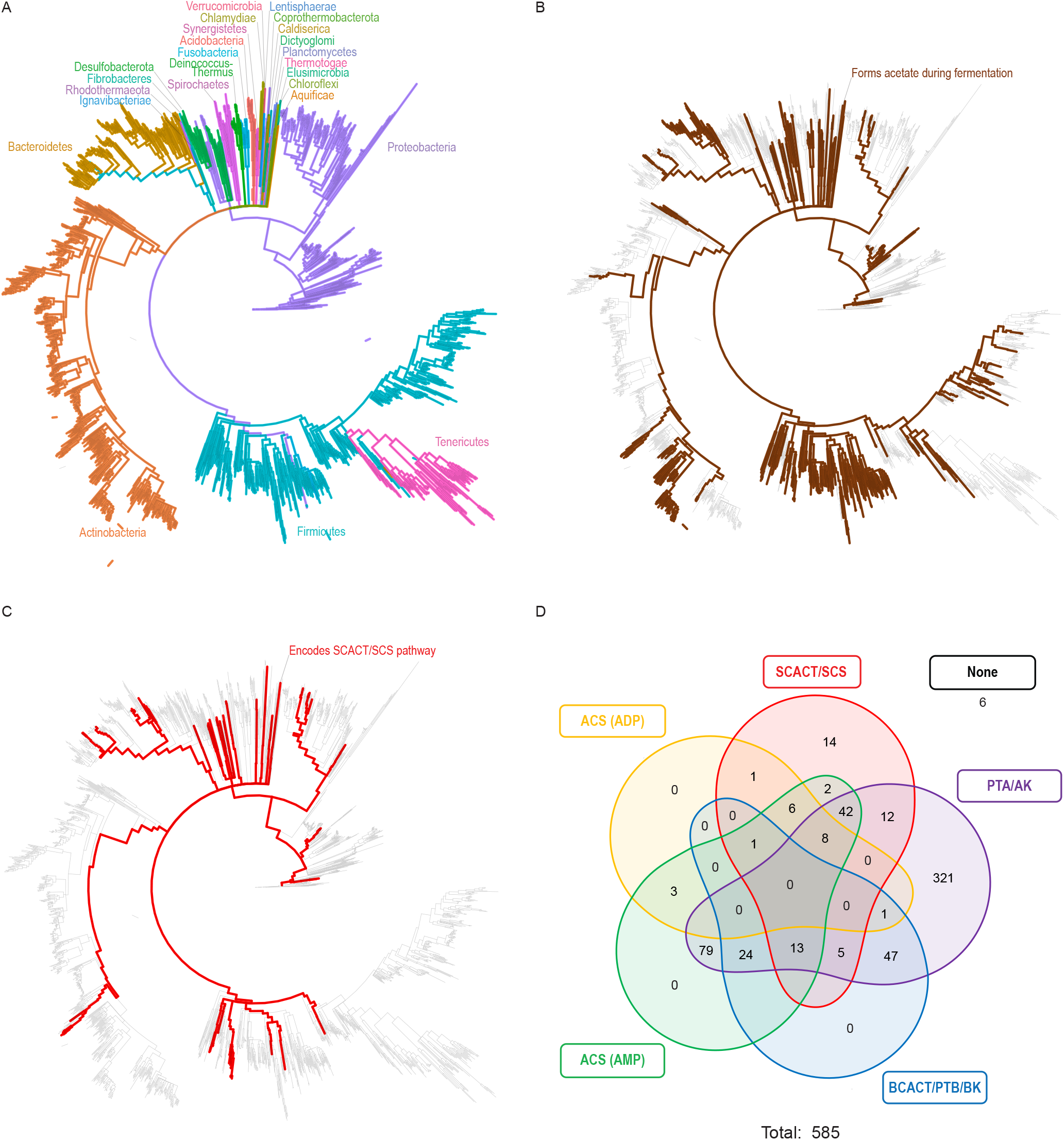
Many bacteria encode the genes of the SCACT/SCS pathway. Phylogenetic tree of type strains of bacteria in Bergey’s Manual, highlighting (A) phyla (B) strains reported to form acetate during fermentation, and (C) strains that encode the SCACT/SCS pathway in their genome. (D) Venn diagram for strains that encode the SCACT/SCS vs. other pathways for forming acetate. In (C) and (D), only strains reported to form acetate during fermentation are included. Abbreviations: ACS (ADP), acetyl-CoA synthetase (ADP-forming); ACS (AMP), acetyl-CoA synthetase; AK, acetate kinase; BCACT, butyryl-CoA:acetate CoA-transferase; BK, butyrate kinase; PTA, phosphotransacetylase; PTB, phosphotransbutyrylase; SCACT, succinyl-CoA:acetate CoA-transferase; and SCS, succinyl-CoA synthetase (ADP-forming).

No archaea were found to encode the SCACT/SCS pathway (Table S1). Instead, all encoded the ACS (ADP) pathway.

## DISCUSSION

We have found a pathway for forming acetate and synthesizing ATP that was previously unrecognized in bacteria. Pathways for forming acetate during fermentation have been studied for over 80 years (6), and so our discovery of the SCACT/SCS pathway is surprising. Nonetheless, it appears to be a common pathway, given it is encoded by nearly 18% of all bacteria that form acetate during fermentation.

We focused on *C*. *granulosum* in this study because it is one bacterium that encodes only the SCACT/SCS pathway. Earlier, we had performed experiments with *Selenomonas ruminantium* HD4, another bacterium that encodes only this pathway (10). In those experiments, we could detect SCS, but not SCACT (11). The reason is unknown but may be because the SCACT had been degraded to an acetyl-CoA hydrolase, an enzyme that we did detect The present experiments confirm that the SCACT/SCS pathway is used in bacteria, even as it leaves the identity of the pathway in *S*. *ruminantium* unknown.

The discovery of the SCACT/SCS pathway in bacteria is important for three reasons. First, it is important for predicting metabolic pathways from genome sequences. These predictions are used to determine the metabolic capabilities of bacteria, especially ones not cultured in the lab. When studies predict metabolic pathways for fermentation, most search for enzymes of the AK/PTA pathway only (11). Some also search for enzymes of the ACS (ADP) pathway (12). The current study shows the importance of looking for enzymes of the SCACT/SCS pathway. Otherwise, studies will overlook pathways (or bacteria) that can form acetate.

Second, it is important to manipulating yield of fermentation products through genetic engineering. One study tried to engineer *A*. *acidipropionici* to produce less acetate (more propionate), but it focused on deleting AK (12). Our study suggests that enzymes of the SCACT/SCS pathway would be more appropriate targets.

Third, this discovery is important to the evolution of metabolic pathways. With this discovery, all known pathways for forming acetate can now be found in bacteria. The SCACT/SCS pathway is not found exclusively in the eukaryotes; if it evolved in eukaryotes, it was an independent event.

## MATERIALS AND METHODS

### Organisms

The DSMZ was the source for all bacterial strains. The strains used were *Acidipropionibacterium acidipropionici* 4, *Butyrivibrio fibrisolvens* D1, *Cutibacterium acnes* DSM 1897, *Cutibacterium granulosum* VPI 0507, and *Propionibacterium freudenreichii* E11.1.

### Media and growth

Strains were grown anaerobically under O_2_-free CO_2_ and with Balch tubes with butyl rubber stoppers, using techniques previously described (13,14). We used PYG medium (DSMZ medium 104). Per liter, PYG medium contained 5 g glucose, 5 g Trypticase peptone (product 211921, BD), 5 g peptone (product 211677, BD), 10 g yeast extract (product 212750, BD), 5 g beef extract (product LP0029, Oxoid), 2.04 g K_2_HPO_4_, 40 mg KH_2_PO_4_, 80 mg NaCl, 20 mg MgSO_4_·7H_2_O, 10 mg CaCl_2_·2H_2_O, 1 mL Tween 80, 5 mg haemin, 1 μL vitamin K_1_, 4 g NaHCO_3_, and 449 mg cysteine-HCl. Resazurin was added as a redox indicator. The haemin was added as a 0.5 g/L solution containing 10 mM NaOH as the diluent. The vitamin K_1_ was added as a 5 mL/L solution containing 95% ethanol as the diluent. The temperature for *A. acidipropionici* 4 and *P. freudenreichii* E11.1 was 30°C. The temperature for *C. acnes* DSM 1897 and *C. granulosum* VPI 0507 was 37°C.

*B. fibrisolvens* D1 was grown on DSMZ medium 712 and at 37°C. Per liter, DSMZ medium 712 contained 9 g glucose, 8.3 g Bacto yeast extract (product 212750, BD), 383.196 mg K_2_HPO_4_, 383.774 mg KH_2_PO_4_, 750.56 mg (NH_4_)_2_SO_4_, 759.72 mg NaCl, 73.426 mg MgSO_4_, 99.967 mg CaC1_2_·2H_2_O, 10.669 g NaHCO_3_, 16.7 mL vitamin solution, and 1.616 g cysteine hydrochloride monohydrate. Per liter, the vitamin solution contained 2 mg biotin, 2 mg folic acid, 10 mg pyridoxine-HCl, 5 mg thiamine-HCl, 5 mg riboflavin, 5 mg nicotinic acid, 5 mg D-Ca-pantothenate, 0.1 mg vitamin B12, 5 mg p-aminobenzoic acid, and 5 mg lipoic acid. Resazurin was added as a redox indicator.

### Cell extracts

Eight, 9-mL cultures were grown to mid-exponential phase and then pooled. Cells were harvested by centrifugation (10,000 *g* for 10 min at 4 °C; F15-8×50cy rotor and Sorvall Legend XTR centrifuge), washed twice in buffer (50 mM Tris [pH = 7.2] and 10 mM MgCl_2_), and resuspended to 3.8 mL in same buffer. All steps after growing the cultures were performed aerobically. The resuspended cells were lysed with a French press (Glen Mills). The resuspended cells were transferred to a mini cell pressure cell and lysed at 16,000 psi. Cell debris was removed by centrifugation (10000 *g* for 10 min at 4°C).

Cell extract was prepared three or more different times for each bacterial strain. It was stored at −80°C until use.

### Supernatant

One 9-mL culture was grown to mid-exponential phase. Cells were removed from supernatant by centrifugation (10,000 *g* for 10 min at 4 °C; F15-8×50cy rotor and Sorvall Legend XTR centrifuge).

Supernatants were prepared three or more different times for each bacterial strain. It was stored at −20°C until use.

### Enzymatic assays

We measured enzymatic activity of cell extracts. Assays were performed at room temperature following conditions in Table 1. Assay products were monitored by measuring absorbance with a Molecular Devices M3 spectrophotometer. Activity was calculated over the time that absorbance increased (or decreased) linearly. Values were corrected by subtracting off the activity of controls (where water replaced cell extract, substrate, or cofactor).

**Table 1.** Conditions used to measure enzymatic activity.

| Enzyme | Reference | Assay components | Product measured (wavelength) | Controls |
| --- | --- | --- | --- | --- |
| Succinyl-CoA:acetate CoA-transferase (SCACT; EC 2.8.3.18) | (30) | 100 mM Tris (pH = 8.0), 0.6 mM succinyl-CoA, 60 mM potassium acetate (pH = 7.0), 0.4 mM L-malate, 1 mM NAD, 2.84 mM KPO<sub>4</sub> (pH = 6.8), 8.8 U/mL malate dehydrogenase<sup>1</sup>, 1.4 U/mL citrate synthase<sup>2</sup> | Reduced NAD (340 nm)<sup>3</sup> | succinyl-CoA, acetate, or cell extract replaced with water |
| Succinyl-CoA synthetase (SCS; EC 6.2.1.5) | (31) | 50 mM Tris (pH = 7.2), 10 mM MgCl<sub>2</sub>, 10 mM succinate (pH = 7.0), 1.2 mM ATP, 0.4 mM CoA, 200 mM hydroxylamine (pH = 7.0) | Succino-hydroxamate (540 nm)<sup>4</sup> | succinate replaced with water |
| Acetyl-CoA synthetase (ACS) (EC 6.2.1.13, EC 6.2.1.1) | (32) (32) | 50 mM Tris (pH = 7.2), 10 mM MgCl<sub>2</sub>, 40 mM potassium acetate (pH = 7.0), 1.2 mM ATP, 0.4 mM CoA, 200 mM hydroxylamine (pH = 7.0) | Aceto-hydroxamate (540 nm)<sup>5</sup> | potassium acetate replaced with water |
| Phosphotransbutyrylase (PTB; EC 2.3.1.19) | After (32) | 76 mM Tris (pH = 7.6), 10 mM NH<sub>4</sub>Cl, 2.5 mM butyryl phosphate, 0.25 mM CoA | Butyryl-CoA (233 nm)<sup>6</sup> | butyryl phosphate replaced with water |
| Butyrate kinase (BK; EC 2.7.2.7) | After (31) | 50 mM Tris (pH = 7.2), 10 mM MgCl<sub>2</sub>, 200 mM butyrate (pH = 7.0), 5 mM ATP, 200 mM hydroxylamine (pH = 7.0) | Butyryl-hydroxamate (540 nm)<sup>5</sup> | butyrate replaced with water |
| Phosphotransacetylase (PTA; EC 2.3.1.8) | (32) | 76 mM Tris (pH = 7.6), 10 mM NH<sub>4</sub>Cl, 5 mM acetyl phosphate, 0.25 mM CoA | Acetyl-CoA (233 nm)<sup>6</sup> | acetyl phosphate replaced with water |
| Acetate kinase (AK; EC 2.7.2.1) | After (31) | 50 mM Tris (pH = 7.2), 10 mM MgCl<sub>2</sub>, 100 mM potassium acetate (pH = 7.0), 5 mM ATP, 200 mM hydroxylamine (pH = 7.0) | Aceto-hydroxamate (540 nm)<sup>5</sup> | potassium acetate replaced with water |

<sup>1</sup>From porcine heart (Sigma M1567-5KU)<sup>2</sup>From porcine heart (Sigma C3260-200UN)<sup>3</sup>Extinction coefficient of 6,220 M<sup>-1</sup> cm<sup>-1</sup><sup>4</sup>Succinic anhydride was used as a standard and prepared in 50 mM Tris (pH = 7.2), 10 mM MgCl<sub>2</sub>, and 200 mM hydroxylamine (33)<sup>5</sup>Acetyl phosphate was used as a standard and prepared in 50 mM Tris (pH = 7.2), 10 mM MgCl<sub>2</sub>, and 200 mM hydroxylamine (pH = 7.0)<sup>6</sup>Extinction coefficient of 4,440 M<sup>-1</sup> cm<sup>-1</sup> (value for S-acetyl-N-succinylcysteamine) (34)

Several assays (for SCS, ACS, BK, AK) measured formation of hydroxamic acids. For these assays, we stopped the reaction at intervals by addition of 0.2857 volumes of development solution (0.25 M FeCl_3_, 2.5 M HCl, and 15% w/v trichoroacetic acid) (15). Afterwards, we held the assay mix for 5 to 30 min at room temperature, centrifuged it at 18,000 *g* for 2 min, and then measured absorbance of the supernatant.

When available, purified enzymes were used as additional controls. Specifically, assays were spiked with purified enzyme at the end of the incubation. The enzymes were obtained from Megazyme. They were SCS from an unspecified prokaryotic source (product code E-SCOAS), ACS (AMP-forming) from *Bacillus subtilis* (product code E-ACSBS), PTA from *B. subtilis* (product code E-PTABS), and AK from an unspecified source (product code K-ACETRM).

One assay measured formation of acetyl-CoA from different combinations of succinate, acetate, ATP, and CoA. The assay contained 100 mM Tris (pH = 8.0), 0.4 mM L-malate, 1 mM NAD, 2.84 mM KPO_4_ (pH = 6.8), 8.8 U/mL malate dehydrogenase, 1.4 U/mL citrate synthase, 30 mM succinate (pH = 7.0), 60 mM potassium acetate (pH = 7.0), 5 mM ATP, and 0.4 mM CoA. Reduced NAD was the product measured, and the amount of acetyl-CoA formed was calculated according to ref. (16).

We determined if activity was different from 0 using an analysis of variance (ANOVA). The statistical model was

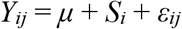

where *Y_ij_* is the observation, *μ* is the overall mean, *S_i_* is the fixed effect of strain, and *ε_ij_* is the residual error. All analyses were conducted in R. The model was fit using the aov function. Least squares means, standard error of the mean, and degrees of freedom were taken from emmeans. *P*-values were calculated using a one-sided *t*-test.

All experiments were replicated at least three times. Each replicate used a different preparation of cell extract.

### Other analyses

Protein concentration in the cell extract was measured using a Pierce BCA protein assay kit (product 23227; Thermo Scientific, Rockford, IL).

Acetate in cell culture supernatant and PYG medium was measured enzymatically. Samples were heated at 100 °C for 10 min to inactivate enzymes. The assay mix contained 150 mM triethanolamine (pH = 8.4), 3 mM MgCl_2_, 10 mM L-malate, 1 mM NAD, 5 mM ATP, 0.4 mM CoA, 1.8 U/mL malate dehydrogenase, 1.4 U/mL citrate synthase, and 0.5 U/mL acetyl-CoA synthetase. Samples were replaced with water as a control. Reduced NAD was measured and acetyl-CoA formed was calculated as above.

We determined if acetate production was different from 0 using the ANOVA model above.

### Analysis of genome sequences

We searched genomes of bacteria and archaea for genes encoding pathways of acetate formation. The genomes were type strains that are 1) reported in Bergey’s Manual (9) and 2) have a genome sequence available on IMG/M (17). The workflow for finding genomes is shown in Fig. 5 and 6. It is similar to a workflow used by us previously (18).

**Figure 5.**
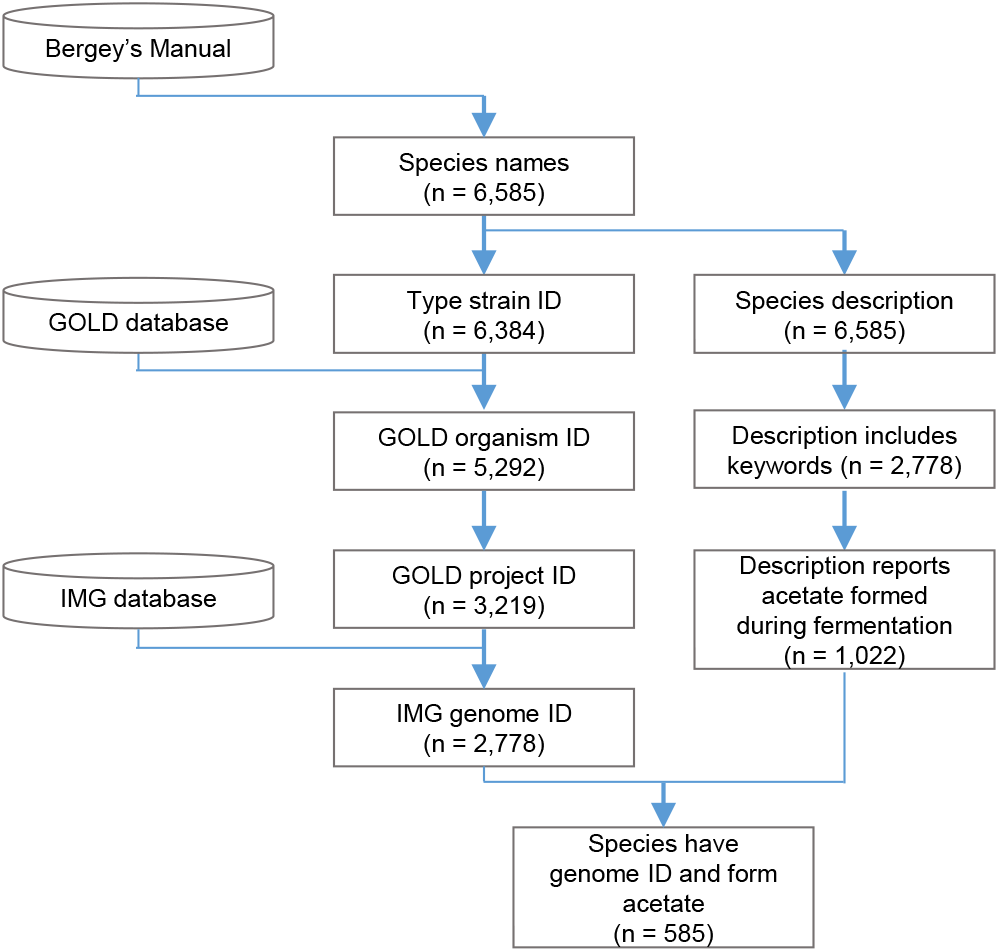
Workflow for finding genomes of type strains of bacteria. Species names and strain IDs were taken from Bergey’s Manual (9). The strain ID was used to find the GOLD organism ID (47), GOLD project ID (47), and the IMG/M genome ID (genome sequence) (17). Though we could have searched IMG/M directly with the strain ID, this approach was slow. Separately, a written species description was searched for keywords (“ferment”, “acetate”, or “acetic”). If keywords were found, the description was read in full to determine if the bacterium forms acetate during fermentation. The description was from Bergey’s Manual, and it included all relevant text from the article for the genus. Our definition of species is broad and includes subspecies, biovars, pathovars, and genomospecies. Our definition of type strains is broad and includes reference or “deposited” strains. Some type strain IDs were generic (e.g., numbers like “238”) and could match multiple GOLD organism IDs. To make matches more specific, we required the species or genus name to match, also.

**Figure 6.**
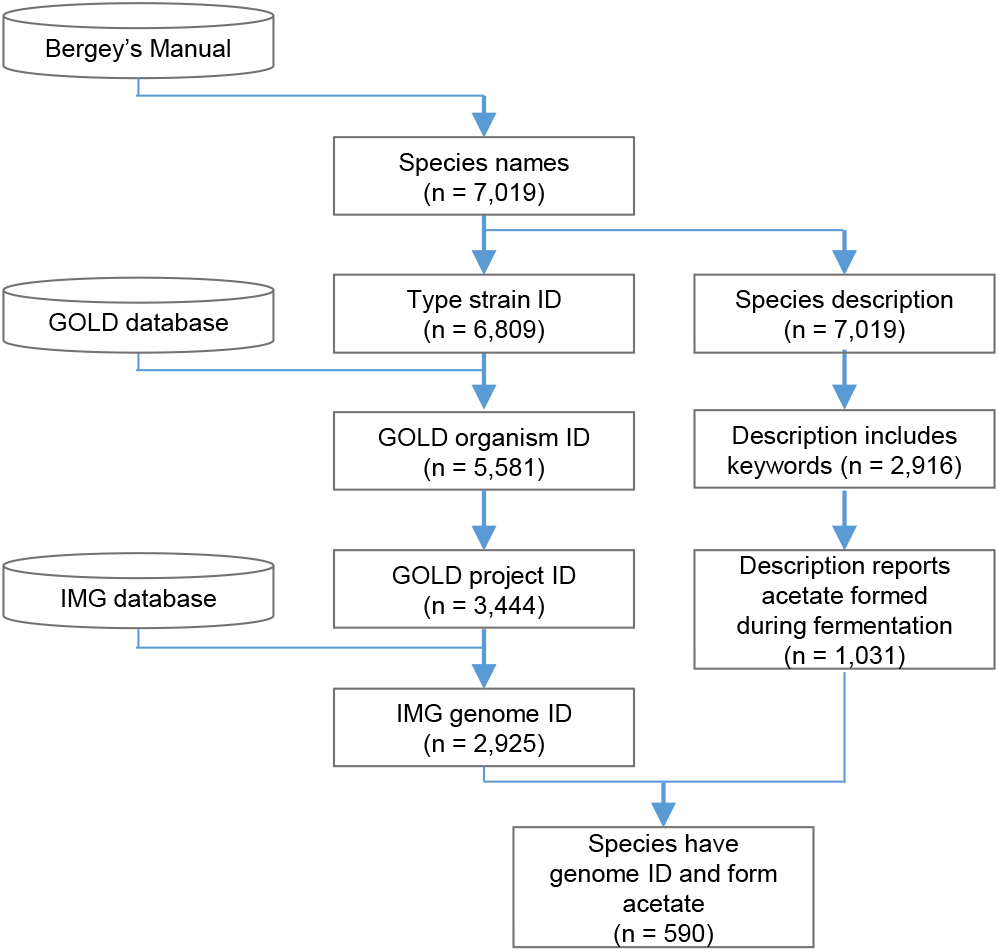
Method for finding genomes of bacteria and archaea that form acetate during fermentation. As Fig. 5, except archaea are included

We searched for genes using IMG/M (17). We searched for each gene’s KEGG Orthology (KO) ID (19) (Table S2). IMG/M sometimes annotated genes with the wrong KO ID (Table S3 to S5). For the genes, we searched with pfam (20) or TIGRFAM (21) IDs instead (Table S2). The locus tags found by these searches are in Table S1.

A pathway was encoded if genes for all enzymes was found. If an enzyme had multiple genes (subunits), all genes for that enzyme had to be found. If a gene had multiple database IDs (domains), all databases ID had to be found, also. If a pathway could be catalyzed by multiple isozymes, genes for only one isozyme had to be found.

### Construction of phylogenetic tree

We constructed a phylogenetic tree of genomes of type strains mentioned above. The construction followed the general approach of ref. (22,23) and used sequences of 14 ribosomal proteins.

We retrieved amino acid sequences of the ribosomal proteins from IMG/M. We retrieved them after first finding the respective genes with KO IDs (Table S6). We discarded sequences that were short (length <75% of the average for a given ribosomal protein). A total of 2,523 genomes had all protein sequences and were analyzed further.

We aligned sequences with Clustal Omega (24,25) and then concatenated them. We discarded columns in the alignment with a large number of gaps (95% or more).

We used aligned and concatenated sequences to create a phylogenetic tree. The tree was calculated using maximum likelihood with RAxML (26) on the CIPRES web server (27). The parameters are listed in Table S7.

Final analysis and visualization were done in R. The consensus tree and branch lengths were calculated using phytools (28). The tree was visualized using ggtree (29). Because no archaea encoded the SCACT/SCS pathway, they were not included in the final visualization of the tree. A total of 2,464 genomes were included in this visualization.

## Notes

### Competing Interest Statement

The authors have declared no competing interest.

